# Scalable co-sequencing of RNA and DNA from individual nuclei

**DOI:** 10.1101/2023.02.09.527940

**Authors:** Timothy R. Olsen, Pranay Talla, Julia Furnari, Jeffrey N. Bruce, Peter Canoll, Shan Zha, Peter A. Sims

## Abstract

The ideal technology for directly investigating the relationship between genotype and phenotype would analyze both RNA and DNA genome-wide and with single-cell resolution. However, existing tools lack the throughput required for comprehensive analysis of complex tumors and tissues. We introduce a highly scalable method for jointly profiling DNA and expression following nucleosome depletion (DEFND-seq). In DEFND-seq, nuclei are nucleosome-depleted, tagmented, and separated into individual droplets for mRNA and genomic DNA barcoding. Once nuclei have been depleted of nucleosomes, subsequent steps can be performed using the widely available 10x Genomics droplet microfluidic technology and commercial kits without experimental modification. We demonstrate the production of high-complexity mRNA and gDNA sequencing libraries from thousands of individual nuclei from both cell lines and archived surgical specimens for associating gene expression phenotypes with both copy number and single nucleotide variants.

## Introduction

In recent years, single-cell barcoding and deep sequencing technologies have conspired to revolutionize our understanding of health and disease by allowing highly-scalable analysis of DNA or RNA with unprecedented resolution^1–3^. Transcriptional states, regulatory networks, and cell state transitions can be identified with scRNA-seq, while mutations (*e.g*., copy number, structural, and single nucleotide variants) can be identified with scDNA-seq. Assessing both the RNA and DNA from thousands of single cells from a single specimen or sample would enable joint identification of rare cell types and subclones from tumors, linkage of cancer driver mutations with distinct expression profiles, and simultaneous analysis of somatic mosaicism and lineage identity in complex human tissues^4^. Such information could be used to understand disease progression, cancer drug resistance, and cell type-specific drug responses, possibly leading to new therapeutic strategies. However, simultaneously assessing the RNA and DNA of individual cells at high throughput remains challenging and crucial. For example, solid tumors might contain many transcriptional subpopulations and genetic subclones in a complex with numerous non-neoplastic cell types in the microenvironment^5, 6^. This challenge can be even more significant in studies of somatic mosaicism in ostensibly normal tissues.

Pioneering single-cell co-assays for RNA and genomic DNA (gDNA), such as gDNA-mRNA-seq (DR-seq)^7^, genome and transcriptome sequencing (G&T seq)^8^, Simul-seq^9^, and simultaneous isolation of genomic DNA and total RNA (SIDR)^10^, have revealed many of the novel insights enabled by combined copy number variation (CNV) and gene expression analysis. These techniques directly linked chromosomal aneuploidies to gene expression and could correlate transcript abundance to copy number variation. They also relied on manually manipulating individual cells, which limited the scale of these techniques to, at most, hundreds-of-cells when performing experiments in multi-well plates. With genotyping of transcriptomes (GoT)^11^, it became possible to detect mutations in the mRNA of targeted loci for thousands of cells with corresponding gene expression information. Other scalable multiomic technology has demonstrated targeted genomic DNA sequencing with targeted surface protein detection^12^. More recently, a scalable combinatorial indexing (sci-) co-assay for RNA and genomic DNA termed sci-L3-RNA/DNA^13^ was introduced. This technique tagments the gDNA of nucleosome-depleted nuclei with barcodes while simultaneously reverse transcribing the RNA to cDNA with a barcoded primer. Additional barcodes are added to the cDNA and gDNA with split-pool ligations. Proof-of-concept studies resulted in co-RNA/DNA libraries for a few thousand cells with very limited coverage, and the experiments were restricted to cell lines.

Here we present DNA and Expression Following Nucleosome Depletion sequencing (DEFND-seq), a scalable method for co-sequencing RNA and DNA from single nuclei that uses commercial droplet microfluidics to achieve a high-throughput. In DEFND-seq, we treat nuclei with lithium diiodosalicylate to disrupt the chromatin and expose genomic DNA (gDNA)^14^. The nuclei are then tagmented, which fragments and tags gDNA with common adapter sequences. Tagmented nuclei are loaded into a microfluidic droplet generator, which co-encapsulates nuclei, beads containing transcriptomic and genomic barcodes, and reverse transcription reagents into single droplets. Ultimately two libraries are created, one for nuclear mRNA and one for gDNA, with each library containing barcodes linked to its originating nucleus, thus allowing simultaneous analysis of the transcriptomes and genomes of individual nuclei. Following nuclear isolation and chromatin disruption, all steps can be performed using a commercial droplet microfluidics system from 10x Genomics and 10x Genomics Chromium Single Cell Multiome ATAC+Gene Expression Kit without further experimental modification.

## Results

Previous reports have demonstrated chromatin disruption in intact nuclei for scDNA-seq^14^. 10x Genomics has commercialized a widely-used droplet microfluidic system for single-nucleus ATAC-seq (snATAC-seq) and joint single-nucleus RNA/ATAC-seq that barcodes thousands of tagmented nuclei with intact chromatin to profile open chromatin. We reasoned that by combining chromatin disruption methods with this commercial system for snATAC-seq, we could instead obtain scDNA-seq data at scale. Furthermore, feeding chromatin-disrupted nuclei to a kit for joint snRNA/ATAC-seq that uses the same commercial platform would yield joint snRNA/DNA-seq data at scale (Figure 1a). Given the large install-base of 10x Genomics droplet microfluidic systems and availability of pre-assembled kits, this advance would make high-throughput, joint profiling of RNA and DNA from individual nuclei broadly available to the research community on a familiar and established platform.

**Figure 1:**
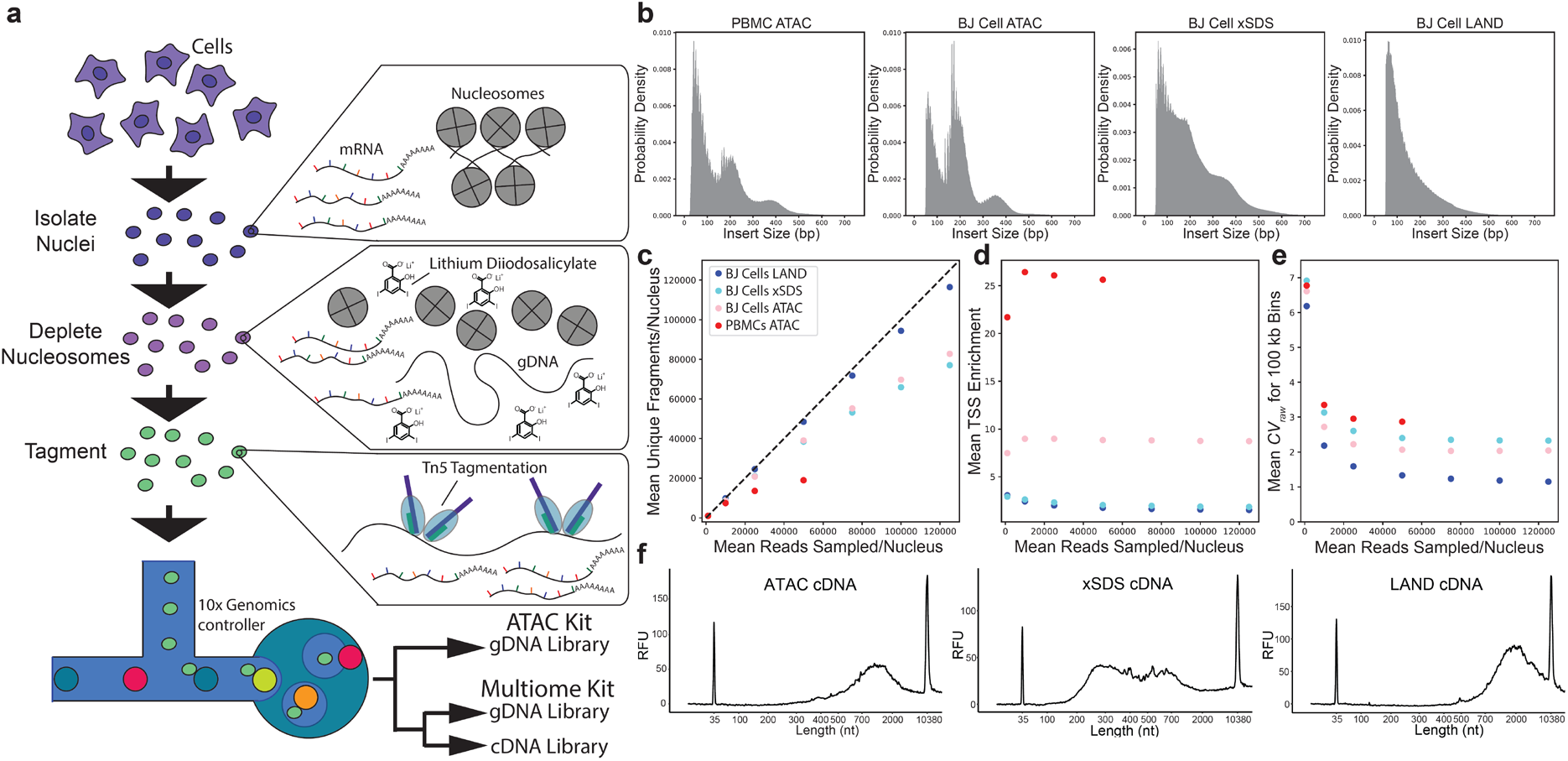
Schmatic and characterization of nucleosome depletion methods. (a) Schematic of the gDNA protocol. Nuclei are isolated, treated with lithium diiodosalicylate, and then tagmented with scATAC-seq reagents or multiome reagents. A gDNA library is produced using scATAC-seq reagents (10x Genomics). When using 10x Genomics Multiome reagents, gDNA and cDNA libraries are produced. We term this DEFND-seq. (b) Fragment length distribution for a scATAC-seq library with PBMCs, scATAC-seq with BJ fibroblasts, BJ fibroblast nuclei prepared using the xSDS protocol, and BJ fibroblast nuclei prepared with the LAND protocol. (c - e) Subsampling analysis of (c) unique fragments, (d) transcription start site (TSS) enrichment, and (e) 100 kb bin coefficient of variation (CV) with BJ fibroblasts prepared with scATAC-seq, BJ fibroblasts prepared with xSDS, BJ fibroblasts prepared with LAND, and PBMCs prepared with scATAC-seq. (f) Bulk BJ fibroblast cDNA BioAnalyzer traces for cells prepared with ATAC, cells prepared with xSDS, and cells prepared with LAND.

We first investigated methods to disrupt chromatin using: (1) cross-linking and sodium dodecyl sulfate-based nucleosome depletion (xSDS), which uses SDS to disrupt chromatin packing with a non-ionic detergent that requires cross-linking to maintain nuclear integrity, and (2) lithium diiodosalicylate assisted nucleosome depletion (LAND), which uses a milder detergent that has previously been used to extract histones^14^. We compared libraries generated from these nucleosome depletion techniques to a standard snATAC-seq library (Figure 1b). Genomic heterogeneity, copy number alterations, polyploidy, and high rates of cell division found in most immortalized cancer cell lines are problematic for characterizing noise and coverage bias. Thus, we chose BJ fibroblasts, a human male euploid fibroblast cell line with diploid somatic chromosomes and a single copy of each sex chromosome, for benchmarking due to their lack of aneuploidy and long doubling time. Nuclei treated with xSDS, LAND, or standard conditions for chromatin preservation were profiled using the 10x Genomics’ Chromium Single-cell ATAC reagents to generate barcoded, tagmented libraries according to the manufacturer’s protocol. Libraries from the snATAC-seq protocol have the characteristic multi-modal fragment length distribution with a high-frequency periodicity indicating the ~10 bp helical pitch of double-stranded DNA and low-frequency features (>100 bp) resulting from periodic binding of nucleosomes to DNA (Figure 1b)^15^. Conversely, the nucleosomal pattern is diminished in libraries from xSDS treated-nuclei and completely absent from libraries from LAND-treated nuclei. The LAND fragment length distribution monotonically decreases from approximately 100-500 bp fragments.

We performed subsampling to determine how read depth impacts various DNA sequencing performance metrics. The number of fragments from LAND-treated nuclei scales almost linearly with sequencing depth out to 125,000 reads/nucleus, while the scATAC-seq and xSDS libraries start to saturate (Figure 1c). This indicates that the LAND libraries are more complex and offer significantly higher coverage of the genome than either xSDS or snATAC-seq libraries, likely because of more complete chromatin disruption. Furthermore, snATAC-seq libraries have transcription start site (TSS) enrichment scores indicating non-uniform coverage of genomic regions due to the presence of nucleosomes, while LAND and xSDS libraries have markedly lower scores (Figure 1d).

To quantify the coverage uniformity of each method, we calculated the raw coefficient of variation (CV) of the number of fragments per 100 kb somatic chromosome bin for several read depths (Figure 1e). Because we are using euploid cells, the CV will ultimately indicate the ability to accurately detect copy number variants (CNVs)^16^. The LAND profiles have the lowest noise (CV ~1.1). Importantly, the CV is still decreasing at higher sequencing depths for the LAND nuclei, while the xSDS and scATAC-seq libraries are saturated.

We also included a similar analysis of a standard benchmarking snATAC-seq dataset^17^ provided by 10x Genomics obtained from peripheral blood mononuclear cells (PBMCs), which are also euploid (Figure 1c—e). This library was of exceptional quality, exhibiting a very high TSS enrichment score (>25) consistent with highly intact chromatin. As expected, this results in low library complexity (saturating with few unique fragments detected per nucleus) and a high CV, indicating low coverage uniformity along the genome, presumably because the tagmented gDNA is stringently limited to open chromatin.

Our ultimate goal is jointly profiling RNA and DNA from the same nuclei. Thus, we also investigated the cDNA yield and quality from the various treatments. We synthesized full-length, bulk cDNA libraries for the different nuclear treatments (Figure 1f). Both the ATAC-seq and LAND nuclei yield full-length cDNA molecules. Interestingly, the xSDS-treated nuclei have truncated cDNA molecules (average length: 786 bp), potentially due to fragmentation or cross-linking.

### DEFND-seq: Joint, Single-Nucleus RNA and DNA Sequencing

Considering that the LAND nuclei exhibited higher library complexity, coverage uniformity, and cDNA quality than alternatives, we further investigated whether this technique could be compatible with combined RNA and DNA sequencing of the same nucleus. We treated nuclei with lithium diiodosalicylate and performed all subsequent steps, without modification, of the 10x Genomics Mulitome kit (see Methods) using BJ fibroblasts and termed this co-assay DNA and Expression Following Nucleosome Depletion Sequencing (DEFND-seq). We obtained DEFND-seq libraries for 1,076 BJ fibroblasts for benchmarking purposes. When compared to the LAND-treated nuclei (which generate a snDNA-seq library without gene expression), the DEFND-seq protocol resulted in no performance degradation with similar TSS enrichment score, CV, and unique fragments (Figure 2a–c). Figure 2d shows the CV distributions for both 100 kb and 1 Mb bin sizes for the full coverage dataset (1.4M reads/nucleus) after correcting for GC and Tn5 insertion bias (see Methods). The DEFND-seq CV (100 kb bins: 0.6, 1 Mb bins: 0.38) is similar to previous reports of other single-cell DNA-seq technologies using 100 kb bins (LIANTI: ~0.2, PTA: ~0.5, AMPL1: ~0.5, QIAGEN MDA: ~0.6, PicoPlex Gold: ~0.6, and GE MDA: 1.0), and 1 Mb bins (LIANTI: 0.1, PTA: ~0.4, AMPL1: ~0.3, QIAGEN MDA: ~0.4, PicoPlex Gold: ~0.6, and GE MDA: 0.5) for BJ fibroblasts, though it must be noted the reported CVs for these methods are using >200× the number of reads/cell than what we report for DEFND-seq and none of these studies demonstrated co-sequencing of RNA and DNA^16, 18^. Thus, the scDNA-seq component of DEFND-seq offers comparable performance to previously published methods for scDNA-seq.

**Figure 2:**
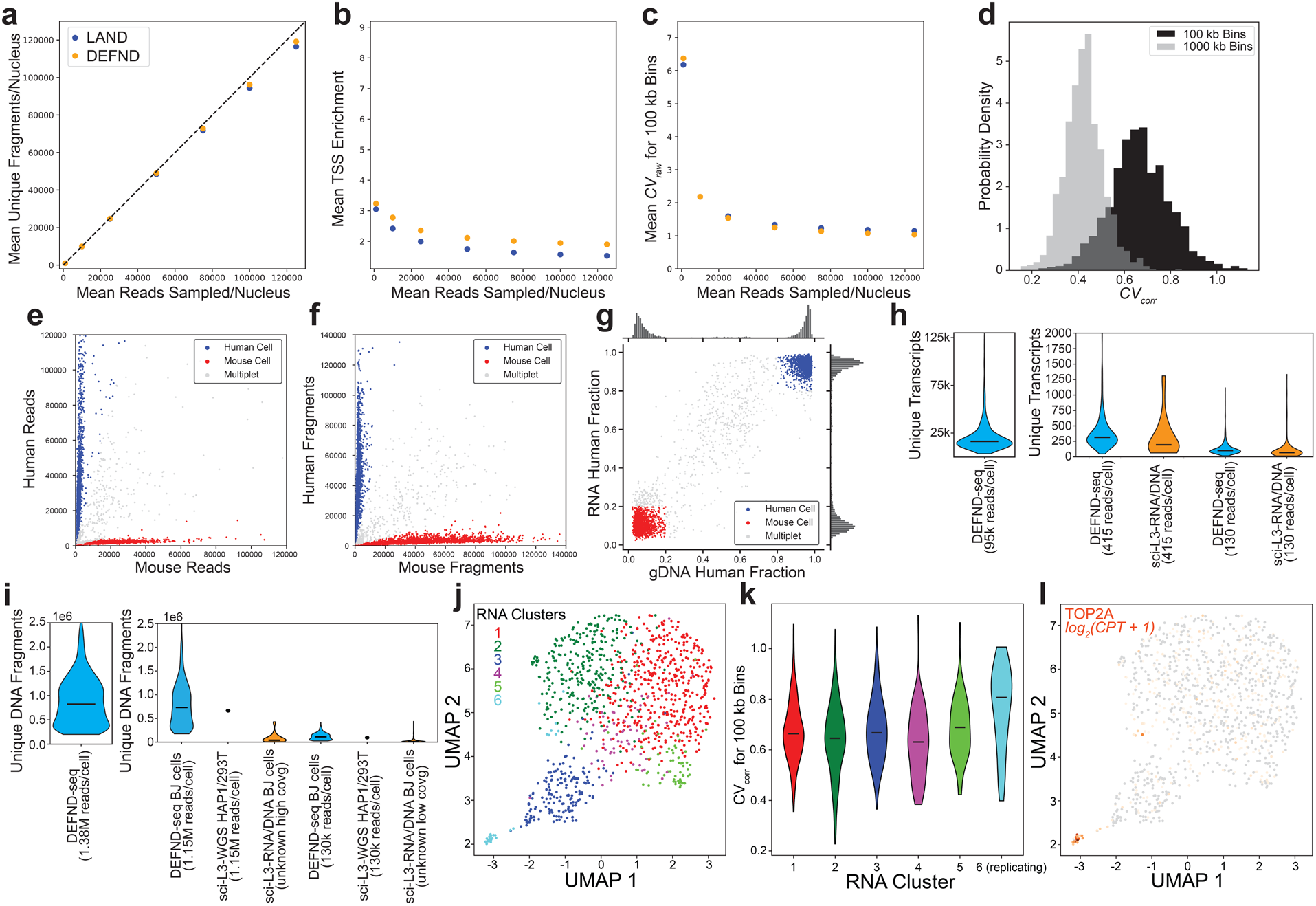
DEFND-seq. (a)-(c) Subsampling analysis of (a) unique fragments, (b) transcription start site (TSS) enrichment, and (c) 100 kb bin coefficient of variation for BJ fibroblasts prepared with LAND and DEFND-seq. (d) Probability density of the coefficient of variation when using 100 kb bins and 1,000 kb bins. (e) Mixed-species gene expression scatterplot. DEFND-seq was performed on a sample with mixed U87 (human) and 3T3 (mouse) cells. Cells were determined to be mouse (>85% of reads are mouse aligned) or human (>80% of reads are human aligned). (f) Mixed-species gDNA scatterplot. Genomic fragments were counted for the same cells in (e). Cells were determined to be mouse (>80% of fragments are mouse aligned) or human (>85% of fragments are human aligned). (g) Scatter plot of the human alignment rates for RNA and gDNA for the mixed species nuclei from (e) and (f). (h) Violin plots of gene expression performance. Bars represent median values. The number of unique transcripts at full sequencing depth is shown, as well as down-sampled violin plots comparing DEFND-seq to sci-L3-RNA/DNA. (i) Violin plots of gDNA performance. Bars represent median values. Total unique gDNA fragments at full sequencing depth are shown along with plots comparing down-sampled DEFND-seq to sci-L3-WGS and sci-L3-RNA/DNA. The sci-L3-WGS data are presented as a single point representing the median. Data for sci-L3-RNA/DNA are of unknown read depth. (j) UMAP embedding colored by gene expression clusters for DEFND-seq BJ fibroblasts. (k) Coefficient of variation of 100 kb genome bins for clusters identified in (j). (i) Same as (j) but colored by TOP2A expression. These cells are primarily located in cluster 6.

To assess the cross-talk and single-nucleus purity of DEFND-seq libraries at high-throughput, we performed a mixed species experiment using human glioma U87 MG cells and murine fibroblast NIH-3T3 cells (Figure 2e–f), profiling 4,400 nuclei in a single 10x Genomics Chromium lane. Inferred human and murine nuclei are represented similarly in both the snRNA-seq (44.8% human, 41.8% murine, and 13.3% mixed) and scDNA-seq data (43.6% human, 44.4% murine, and 11.9% mixed). The mitochondrial content of these cells was relatively low, <5% for all cell types (Supplementary Information, Figure S1). Inferred singlets were quite pure for both the RNA libraries (median purity: 94.1% for inferred human nuclei, 90.3% for inferred murine nuclei) and DNA libraries (median purity: 96.0% for inferred human nuclei, 93.9% for inferred murine nuclei). The multiplet rate aligns with microscopic images of the nuclei taken before tagmentation (Supplementary Information, Figure S2). The multiplet rate will depend on cell type, cell concentration during nuclear isolation and loading density, and can be further reduced by implementing flow sorting, which was not performed for any experiments herein. Moreover, the gene expression and gDNA analyses were concordant; cells that have a high fraction of human reads in the gene expression data likewise have a high fraction of human reads in the genomic DNA data (Figure 2g).

We wanted to compare DEFND-seq to other high-throughput single-cell (thousands of cells per sample without complex automation) DNA/RNA co-assays. The previously reported sci-L3 technique, in principle, fits these criteria^13^. Instead of relying on droplet microfluidics to create nanoliter-sized reactions, in sci-L3 cells undergo xSDS treatment and then are processed through a series of split-pool barcoding reactions, which add unique cell-specific barcodes to the genomic DNA. It has two embodiments: a co-assay that concurrently adds RNA barcodes and DNA barcodes to individual nuclei termed sci-l3-RNA/DNA, and a whole genome version that does not have gene expression capabilities termed sci-L3-WGS. Yin et al. analyze BJ fibroblasts and report median transcripts per nucleus at two very low sequencing depths (415 reads/cell and 130 reads/cell). Although our BJ GEX dataset has ~20,000 median unique transcripts at full depth, we substantially downsampled our data to their reported read depths and found DEFND-seq to detect more unique transcripts using 415 reads/cell (DEFND-seq: 316 median unique transcripts, sci-L3-RNA/DNA: 194 median unique transcripts) and using 130 reads/cell (DEFND-seq: 101 median unique transcripts, sci-L3-RNA/DNA: 68 median unique transcripts) (Figure 2h). Thus, sci-L3-RNA/DNA may have relatively low RNA capture efficiency. Ideally, we would compare our corresponding gDNA dataset to the corresponding sci-L3-RNA/DNA gDNA dataset; however, the sequencing depths are unreported for the gDNA libraries. Instead, we compare DEFND-seq gDNA to the sci-L3-WGS assay, which uses the same chemistry as sci-L3-RNA/DNA to create a gDNA library but does not generate a gene expression library (Figure 3i). This assay reports median unique fragments per nucleus for given sequencing depths, but, unfortunately, uses a polyploid cell line (HEK293T) different from our study. Nevertheless, we downsampled the DEFND-seq data to the reported depths in sci-L3-WGS and found DEFND-seq to have more median unique fragments than sci-L3-WGS at 1.15M reads (DEFND-seq: 731,814 median unique fragments, sci-L3-WGS: 660,700 median unique fragments) and 130k reads (DEFND-seq: 114,314 median unique fragments, sci-L3-WGS: 97,300 median unique fragments). For reference, we note that the sci-L3-RNA/DNA assay reports median unique fragments per nucleus for BJ fibroblasts at two unknown, but likely low sequencing depths (sci-L3-RNA/DNA high: 40,681 median unique fragments, sci-L3-RNA/DNA low: 12,118 median unique fragments). Thus, we conclude that DEFND-seq performs comparably to, if not significantly better than, competing scalable methods for joint single-nucleus RNA/DNA profiling but with the key advantages of a widely used commercial platform and far more extensive benchmarking (Supplementary Information, Figure S3-S4).

**Figure 3:**
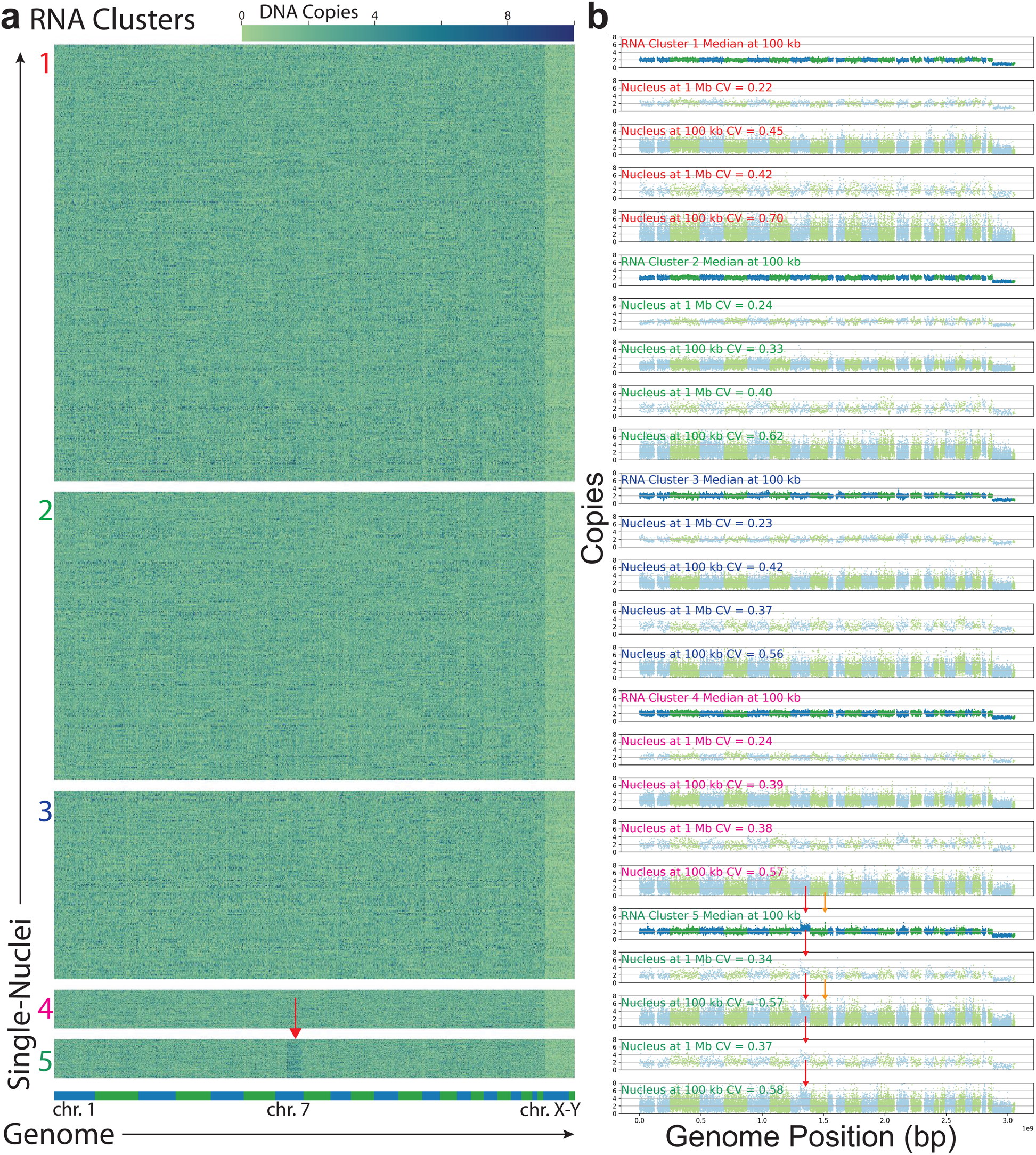
Copy number variation from DEFND-seq analysis of BJ fibroblasts. (a) CNV heat maps with 100 kb bins of individual nuclei grouped by gene expression cluster number (1 – 5). The actively replicating cluster (cluster 6) was omitted because of the aberrant genomes of replicating cells. A chromosomal map colored by chromosome is included as a reference on the bottom margin. Red arrow indicates a partial Chr. 7q amplification in cluster 5. (b) Relative CNV plots for clusters in (a). For each cluster a plot is included for: cluster-median CNV with 100 kb bins, a single low-CV nucleus CNV with 1 Mb bins, the same low-noise nucleus CNV with 100 kb bins, a randomly selected nucleus with near-median CV using 1 Mb bins, and the same near-median CV nucleus with 100 kb bins. Labels are colored by cluster. Bin colors alternate for different chromosomes. Red arrows and orange arrows indicate the Chr. 7q amplification found in cluster 5 and the HAS2 focal amplification in Chr. 8, respectively.

DEFND-seq allows us to directly link mRNA phenotype to genotype. We used the BJ fibroblast gene expression dataset for unsupervised phenotypic clustering of individual nuclei (Figure 2j). Then, using the gDNA dataset, we analyzed the coverage uniformity (CV) of each gene expression cluster (Figure 2k). As expected, actively replicating cells identified from the gene expression data have lower coverage uniformity (higher CV) due to on-going DNA replication (Figure 2l), as observed in previous studies^16^.

Copy number variation (CNV) heat maps and fragment counts for all cells were generated and grouped by gene expression cluster (Figure 3a). We observe uniform coverage of somatic chromosomes in clusters 1-4 and an expected decrease in copy number for the sex chromosomes (male-derived cells). Individual nuclei for a given cluster can be consolidated in low-noise, RNA cluster-level, pseudo-bulk CNV analysis (Figure 3b). For each gene expression cluster corresponding to non-cycling cells, we also present the CNV plots of a single low-noise nucleus with 1 Mb bins, a single low-noise nucleus with 100 kb bins. A randomly selected nucleus with near-median CV is also plotted with 100 kb bins and 1 Mb bins for each cluster (Figure 3b). Interestingly, despite using a healthy euploid cell line, we observe a cluster with an amplification in part of Chr. 7q. This amplification is evident at the cluster level and in individual nuclei (Figure 3b). We also observed focal amplification of *HAS2*. The cells that bear these amplifications are rare (~2%) but the genomic alterations have transcriptional consequences. Indeed, the RNA cluster that is enriched in nuclei harboring partial amplification of 7q are strongly enriched in the expression of genes in the amplified region based on GSEA (Supplementary Information, Figure S5). Thus, even in these benchmarking experiments, we demonstrate that DEFEND-seq, can identify rare phenotypic cellular subpopulations that are enriched in rare genetic subclones.

### DEFND-seq analysis of CNVs, SNVs, and gene expression in a cryopreserved glioblastoma surgical specimen

Glioblastoma (GBM) is an aggressive primary brain tumor with a median survival of 15 months^19^. GBMs are noted for their high degree of phenotypic plasticity and heterogeneity, which complicates pharmacological intervention. Large-scale exome and genome sequencing studies have identified multiple^20, 21^ key driver mutations of GBM, including amplifications and mutations in growth factor receptors (*PDGFRA, EGFR*), *TP53* mutations, and mutations in the PI3K signaling pathway, including *PTEN*. However, relatively few genetic alterations have been associated with established phenotypic subtypes of GBM. Examples include mutations in IDH1 and amplification of *PDGFRA* with the proneural phenotype and NF1 mutations with the mesenchymal subtype^22^. scRNA-seq analysis of GBM has shown that transformed glioma cells can take on any of multiple phenotypic states with varying degrees of neural lineage resemblance and that these states recur across patients^23^. Furthermore, these transcriptional states appear to be highly plastic^24, 25^, which may result in a complicated relationship between somatic mutations and transcriptional states. In addition, recent studies in acute slice cultures^26^ have uncovered cell type-specific drug responses in GBM, but have relied on transcriptional profiling to define drug-sensitive cellular subpopulations, which could be further refined by their genetics. Thus, joint single-cell RNA/DNA sequencing of individual nuclei could be a compelling approach to analyzing these tumors and their response to therapy.

We applied DEFND-seq to a biobanked, cryopreserved primary IDH1 wildtype glioblastoma resection from which we had obtained and reported scRNA-seq profiles of corresponding fresh tissue several years ago^23^. Our previous work showed that this tumor’s transformed cells exhibited exceptional diversity, including neural progenitor-like cells, astrocyte-like cells, cycling cells, mesenchymal-like cells, and oligodendrocyte progenitor-like cells. In addition, there is an unusually small population of non-neoplastic cells (mainly oligodendrocytes), which we expect to be euploid. Thus, detecting and identifying these cells requires high-throughput and sensitivity. Furthermore, unlike when the sample was initially sequenced using fresh tumor tissue and the mRNA from whole cells, the sample has been cryopreserved for >4 years, and cDNA libraries would need to be constructed from nuclear mRNA. Thus, this previously characterized tumor presents multiple compelling challenges for demonstrating the utility and broad applicability of DEFND-seq.

DEFND-seq on the GBM tumor yielded a gene expression and gDNA dataset including 1,821 nuclei with a median of 4,421 transcripts and ~214,000 unique DNA fragments detected per nucleus (at ~360,000 DNA reads/nucleus). Clustering the gene expression data and performing differential gene analysis on the clusters identifies a small cluster of non-neoplastic oligodendrocytes expressing myelin-associated genes, including *MOBP* (33 cells) (Figure 4a—b).

**Figure 4:**
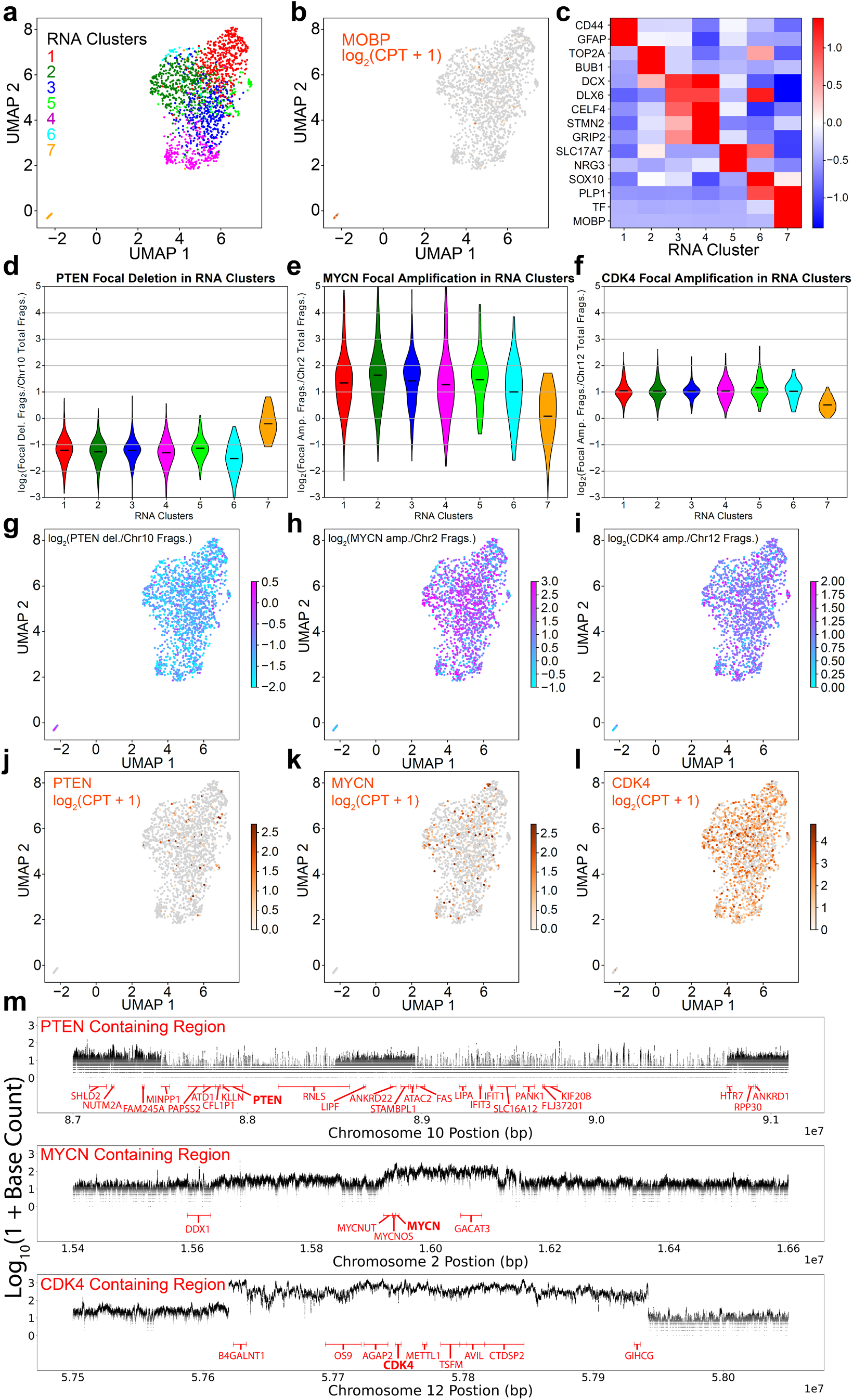
DEFND-seq applied to cryopreserved GBM tumor sample. (a) UMAP embedding of gene expression data colored by unsupervised clustering. (b) Same as (a) but colored by expression of *MOBP*. (c) Differentially expressed genes for each cluster in (a). (d) Single-cell distributions of the log-ratio of the average unique fragements/100 kb bin detected in a focally deleted region containing *PTEN* to the average unique fragments/100 kb bin for Chr. 10 for each cluster in (a) showing focal deletion in clusters 1-6. Bars indicate the medians of each cluster. (e) Same as (d) but for a focally amplified region containing *MYCN* relative to Chr. 2 showing focal amplification in clusters 1-6. (f) Same as (e) but for a focally amplified region containing *CDK4* relative to Chr. 12 showing amplification in clusters 1-6. (g) – (i) Nuclei from (a) colored by genomic log fold change in the CNV-containing regions corresponding to (g) *PTEN*, (h) *MYCN*, and (i) *CDK4*. (j) – (l) Nuclei from (a) colored by mRNA expression (log counts-per-thousand) for (j) *PTEN*, (k) *MYCN*, and (l) *CDK4*. (m) Read count pileup of every base along selected genomic regions for all transformed cells. *PTEN*, *MYCN*, and *CDK4* regions are presented along with a selection of genes located in the sampled window.

We also identified six clusters of putatively transformed cells with the expected neural progenitor-like (*DCX, STMN2*), astrocyte-like (*CD44, GFAP*), proliferative (*TOP2A, BUB1B*), and oligodendrocyte progenitor-like (*SOX10, PLP1*) phenotypes along with a transformed cluster resembling neurons (*SLC17A7, NRG3*) as we reported previously for this tumor^23^ (Figure 4c). The transformed clusters contain a focal deletion of a region of Chr. 10 containing *PTEN*, which is not present in the non-neoplastic oligodendrocyte cluster (Figure 4d, g, j). Deletions of *PTEN* are common in GBM and associated with shorter survival^27^. In this tumor, it appears there has been biallelic deletion of *PTEN*, as one copy of Chr. 10 is lost and the other harbors this focal deletion. The deleted region, which includes *PTEN* has a complex pattern with alternating ~1 Mb-sized focal deletions (Figure 4m).

The transformed clusters also harbor an amplification of a region in Chr. 2 that includes the oncogene *MYCN* and a region in Chr. 12 containing *CDK4* (Figure 4e-f). Amplification of both genes is associated with GBM tumorigenesis^28^. While the transcription factor *MYCN* is not highly expressed, the corresponding CNV is present throughout the transformed clusters (Figure 4h, k). Furthermore, *CDK4*, which is normally expressed during the G1-to-S transition of the cell cycle, is constitutively and highly expressed throughout the transformed clusters (Figure 4i, l). Other highly recurrent aneuploidies such as Chr. 7 amplification and chromosome 10 deletion, which are frequently used to identify transformed glioma cells in single-cell genomics^23^, are directly detected in the transformed cells, as is a complex series of amplified and deleted regions in Chr. 3 (Extended Data Figure 1). Many of these single-cell CNAs are occurring at the sub-megabase level (*e.g*., *MYCN* amplification: ~200 kb long), which would have been obfuscated with larger bins (Figure 4m).

The scalability of genomic profiling technologies like DEFND-seq are ultimately limited by sequencing costs, particularly the DNA sequencing component. Fortunately, sequencing costs are declining, and new entrants to the sequencing market may offer compelling advantages in terms of price, accuracy, and speed. While all of the DEFND-seq data shown so far has been generated using Illumina sequencers, we also tested DEFND-seq on the recently commercialized Element Aviti, resequencing the GBM libraries described above. Subsampling analysis revealed that the Element Aviti dataset had similar library complexity but with reduced transcription start site enrichment and ~25% longer fragments when compared to the same library sequenced with an Illumina NovaSeq 6000 (Figure 5 a—c), consistent with a less biased sampling of the library by the Element Aviti sequencer.

**Figure 5:**
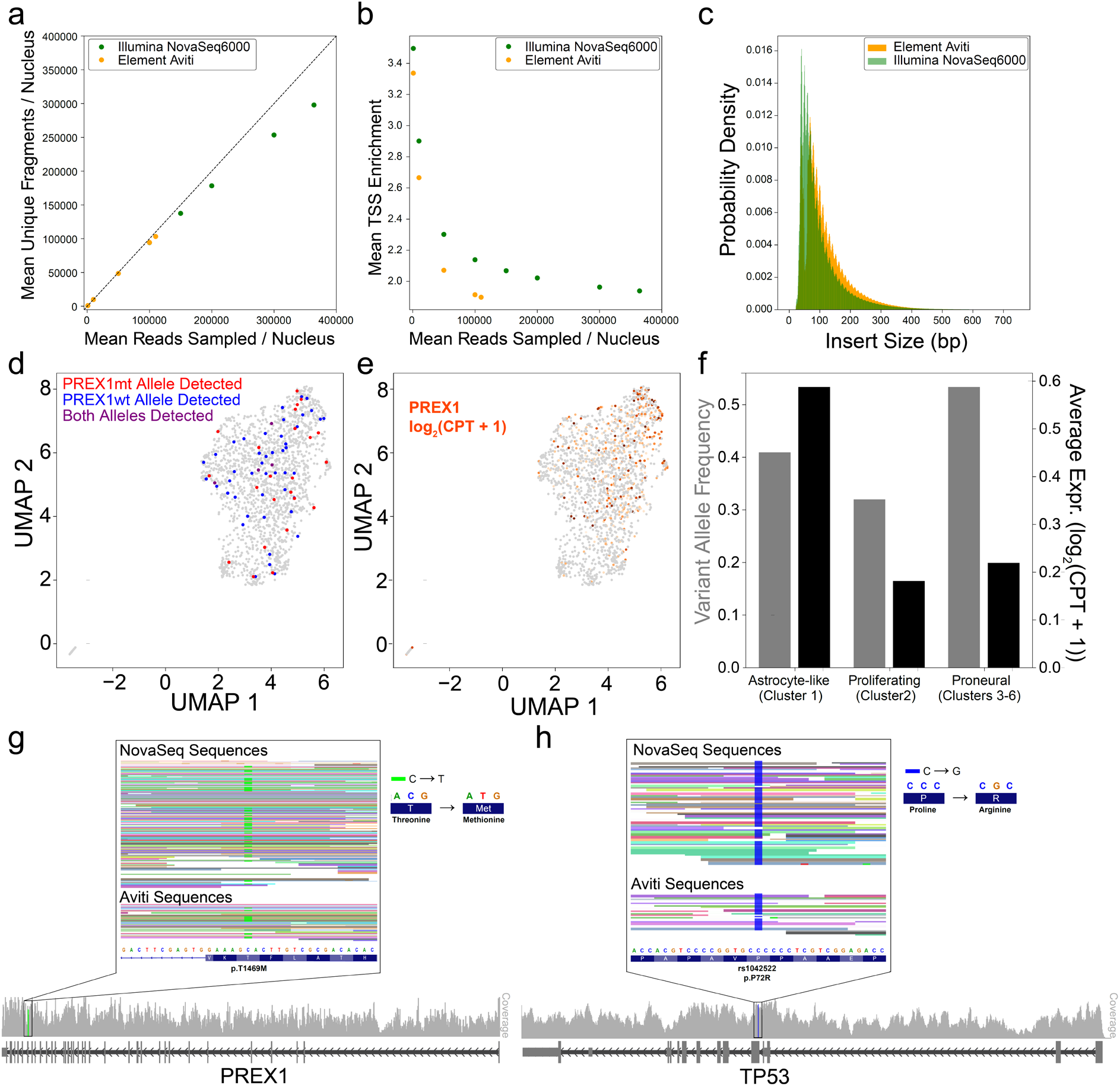
SNP and SNV detection. (a) Sequencing saturation analysis of the number of unique fragments per nucleus for a GBM DEFND-seq library sequenced on an Illumina NovaSeq 6000, and on an Element Aviti. (b) Sequencing saturation analysis of the transcription start site enrichment for a library sequenced on both Illumina NovaSeq 6000 and Element Aviti sequencers. (c) Insert size distribution for the GBM library sequenced on Illumina NovaSeq 6000 and Element Aviti sequencers. (d) *PREX1* genomic mutation detection projected on a UMAP derived from the gene expression dataset. (e) Same as (d) but colored by *PREX1* expression. (f) *PREX1* variant allele frequency and average expression of *PREX1* in astrocyte-like, proliferating, and proneural clusters. (g) Integrated genomic viewer (IGV) style plots of the transformed cells showing exonic regions of *PREX1*, coverage, and the location of the SNV^63^. The SNV is magnified to show the missense mutation and affected amino acid. Individual reads are shown for both the Illumina NovaSeq 6000 and Element Aviti sequencers and each read is colored by cell of origin (cell barcode). (h) Same as (g) but for the p.P72R mutation on *TP53*.

We were interested in DEFND-seq’s ability to analyze single nucleotide polymorphisms and somatic variants (SNPs and SNVs). Furthermore, having sequenced the same library with two different sequencers could provide increased accuracy since the two technologies presumably have different error profiles. Unfortunately, the material for sequencing the germline of this patient was unavailable. Thus, we identified likely somatic SNVs by intersecting the mutation calls from the two sequencers and filtering on large databases of common germline SNPs (see Methods). These putative mutations were then further filtered against the catalogue of somatic mutations in cancer (COSMIC)^29^, leading to the identification of a missense point mutation in *PREX1* (p.T1469M) (Figure 5g). *PREX1* encodes a guanine exchange factor and is part of the PI3K signaling pathway, which is often mutated and activated in GBM. We detected either the wildtype, mutant, or both alleles in 76 transformed cells with an overall variant allele frequency of 0.38, suggesting that it is subclonal (Figure 5d). As we might expect for highly plastic transcriptional states in GBM, there is no statistically significant bias in the representation of the *PREX1* mutant allele across clusters (Figure 5e). However, we found that *PREX1* expression is significantly enriched among the astrocyte-like glioma cells (cluster 1, ~2.2-fold enriched, FDR<0.00001) (Figure 5f). Thus, DEFND-seq can discern relationships between genotype and phenotype for both somatic SNVs and CNVs.

Although we did not find common hotspot somatic mutations in *TP53* in this tumor, the patient harbors a p.P72R variant in *TP53* (Figure 5h). While this variant has been confirmed to be somatic in several tumor types, including one GBM case in COSMIC^30^, it also corresponds to a well-known common SNP (rs1042522). Cells containing the p.P72R variant reportedly have decreased inhibition of PGC-1α, which is a master regulator of mitochondrial biogenesis and oxidative phosphorylation. This may increase the migration and invasion capacity of tumor cells since these p.P72R cells have greater mitochondrial function^31^. Furthermore, p53 with the p.P72R mutation has a greater ability to bind to p73 and neutralize p73-induced apoptosis^32^. All reads in every cell (53 cells total) with coverage of this locus contain this variant, implying homozygosity. Nonetheless, previous studies using The Cancer Genome Atlas (TCGA) have shown that heterozygous individuals with GBM exhibit loss of heterozygosity in their tumor tissue in favor of the p.P72R allele^33^. However, we do not observe a deletion at this locus in our DEFND-seq data, and conclude that this is likely a homozygous SNP.

## Discussion

We have described a method for scalable co-sequencing of RNA and DNA from individual nuclei on a broadly available platform using commercial reagents. Nuclei are depleted of their chromatin packing through exposure to lithium diiodosalicylate, which sufficiently maintains the structure of the nucleus to allow manipulation of individual nuclei and extraction of high-quality mRNA with a droplet generator. Compared to libraries prepared without chromatin depletion or with other chromatin depletion methods, the LAND-treated nuclei yield higher complexity libraries and more uniform genomic coverage. Furthermore, we can generate full-length cDNA libraries from LAND-treated nuclei. Compared to Yin et al.’s co-assay for DNA and RNA, we demonstrated much higher performance with extensive benchmarking^13^. Additionally, DEFND-seq is implemented on a platform that is widely and routinely used in single-cell studies without modification of the equipment or protocols. We performed experiments with up to 4,400 nuclei per chip lane but could readily scale this to ~40,000 nuclei per sample given the 8-lane capacity of a standard 10x Genomics chip and potentially >100,000 nuclei per flow cell with standard super-loading and demultiplexing with either SNP genotypes^34, 35^ or nuclear hashtag antibodies^36^.

By applying DEFND-seq to a cryopreserved GBM tumor sample we were able to identify rare cells and investigate genotype-phenotype relationships in this complex tumor, including CNVs and SNVs. The snRNA-seq data obtained with DEFND-seq recapitulated our previous observations from whole-cell scRNA-seq on a fresh specimen from the same patient^23^. Most importantly, this experiment demonstrated that high-quality DEFND-seq data can be obtained from archival human tumor tissue stored for several years, suggesting broad applicability to prospective and retrospective clinical studies of human disease.

For simplicity, we implemented DEFND-seq without modification to the 10x Genomics Multiome protocol, but adjusting some steps may lead to more optimal results. For example, DEFND-seq libraries tend to have many more fragments than scATAC-seq libraries, thus reducing the number of polymerase chain reaction cycles may be beneficial for deeply sequenced libraries. Furthermore, some nuclei may be too delicate to sustain the lithium treatment and may require a lower concentration of lithium diiodosalisylate for use with DEFND-seq. In this situation, some optimization may be needed to ensure efficient nucleosome depletion. The results presented here were obtained with minimal optimization, so there is likely room for significant performance improvements.

We anticipate that DEFND-seq can be combined with other multi-modal readouts for single-cell profiling. For example, by combining DEFND-seq with multiplexed targeted amplification either on-chip or retrospectively following pooled library construction on either the mRNA or gDNA libraries, we could likely enhance our genotyping capacity significantly for specific, targeted loci^11, 37^. DEFND-seq should be seamlessly compatible with CRISPR-based pooled screening^38, 39^ and CRISPR^40, 41^ or static barcode-based^42^ lineage tracing. Taken together with the scalability of DEFND-seq, we expect broad applicability to basic and clinical questions from somatic mosaicism to tumor evolution and drug resistance.

## Methods

### Cell Culture

3T3 cells (ATCC #CRL-1658) and U87 MG cells (ATCC #HTB-14) were cultured in Dulbecco’s Modified Eagle’s Medium (ATCC #30-2002) supplemented with 10% feline bovine serum (FBS, ThermoFisher # A3160401). Eagle’s Minimum Essential Medium (ATCC #30-2003) was used to culture BJ cells (ATCC CRL-2522). TrypLE Express Enzyme (ThermoFisher #12605010) was used for dissociating cells during cell passages.

### scATAC-seq

Nuclei were prepared from cryopreserved BJ fibroblasts. Frozen cells were thawed and added to a Dounce homogenizer containing 1 mL of nuclei isolation buffer (NIB: 10 mM TrisHCl pH 7.4, 10 mM NaCl, 3 mM MgCl_2_, 0.1% Igepal (Millipore Sigma #18896), and 1 x protease inhibitors (Millipore Sigma #118735800001)). The sample was gently homogenized on ice with 5 strokes of the loose pestle and 5 strokes of the tight pestle. The cells were transferred to a conical and 5 mL of PBS (Gibco #10010-031) was added. The sample was spun down at 500 g for 5 minutes, washed with 10 mL of PBS and resuspended in 100 μL of 10x Genomics Nuclei Buffer (10x Genomics, #2000207). Single-cell ATAC libraries were prepared with these nuclei using the 10x Genomics Single Cell ATAC kit (10x Genomics #1000176, #1000162, and #1000212) according to the manufacturer’s protocol.

### xSDS Preparation

Nuclei that were cross-linked and then treated with sodium dodecyl sulfate (SDS) to remove nucleosomes (xSDS) were prepared following Vitak et al.^14^ Briefly, BJ fibroblasts were washed with 10 mL of PBS and covered in 3 mL of TrypLE (ThermoFisher #12605028) for 10 minutes at 37 °C. Then 9 mL of media was added, quenching the dissociation, and cells were spun down. The cells were resuspended in 10 mL of media with 406 μL of 37% formaldehyde (Millipore Sigma #252549) and gently rotated at room temperature for 10 minutes. 2.5 M glycine (800 μL) was then added and the suspension was incubated on ice for 5 minutes. Cells were centrifuged at 500 g for 8 minutes and washed with 10 mL of PBS. 5 mL of NIB was added to the pellet and the suspension was incubated for 20 minutes at 4 °C under rotation. Cells were then spun down at 500 g for 5 minutes and washed with 900 μL of NEBuffer 2.1 (New England Biolabs #B7202). The cells were pelleted (500 g for 5 minutes) and resuspended in NEBuffer 2.1 with 12 μL of 20% SDS. The solution was placed in a gentleMACS and vigorously shaken at 42 °C for 30 minutes. 200 μL of 10 % Triton-X (Millipore Sigma #T8787) was added and the cells were again shaken at 42 °C for 30 minutes. The cells were pelleted (500 g for 5 minutes) and resuspended in 100 μL of 10x Genomics Nuclei Buffer, counted, and tagmented following the Chromium Single Cell ATAC protocol. Tagmented nuclei were loaded onto a 10x Chromoium controller and libraries were prepared according to the Chromimum Single Cell ATAC protocol (10x Genomics #1000176, #1000162, and #1000212).

### LAND Preparation for BJ Fibroblasts

The LAND protocol for adherent cells using 10x reagents was adapted from the Vitak et al. LAND protocol^14^. BJ fibroblasts were first washed with 10 mL of PBS and dissociated with 3 mL of TrypLE for 10 minutes at 37 C. Media (9 mL) was used to quench the dissociation reaction and collect the cells. The cells were spun down (300 g for 5 minutes) and 200 μL of DEFND buffer (175 μL NIB, 10 μL 1 mg/mL protease inhibitor (Millipore Sigma #11429868001), 25 μL 100 mM lithium diiodosalicylate (Millipore Sigma #653-14-5)), was added and incubated on ice for 5 minutes. Immediately following this incubation, 10 mL of nuclei isolation buffer was added and the nuclei were centrifuged at 4 °C for 5 minutes at 500 g. The supernatant was removed and 100 μL of 10x Genomics Nuclei Buffer. A fraction of nuclei was stained with SYBR green and counted on a Countess with a GFP filter set. Nuclei were then tagmented and prepared for single-cell sequencing with the Chromium Single Cell ATAC kit (10x Genomics #1000176, #1000162, and #1000212) and protocol.

### DEFND-seq for BJ Fibroblasts

The DEFND protocol followed the LAND protocol with the addition of RNAse inhibitors. Briefly, BJ fibroblasts were washed and dissociated with TrypLE. The cells were spun down and 200 μL of DEFND buffer with RNAse inhibitors (170 μL NIB, 10 μL 1 mg/mL protease inhibitor, 25 μL 100 mM lithium diiodosalicylate, 5 μL SUPERase In (ThermoFisher #AM2694)), was added and incubated on ice for 5 minutes. Immediately following this incubation, 10 mL of nuclei isolation buffer with 10 μL SUPERase In was added and the nuclei were centrifuged at 4 °C for 5 minutes at 500 g. The supernatant was removed and 100 μL of 10x Genomics Nuclei Buffer, with suggested RNAse inhibitors for multiome experiments. A fraction of nuclei was stained with SYBR green and counted on a Countess with a GFP filter set. Nuclei were then tagmented and prepared for single-cell sequencing with the Chromium Single Cell Multiome kit (10x Genomics #1000285, #1000230, #1000215).

### DEFND-seq for Cryopreserved Tissue

A cryopreserved IDH wildtype Grade IV GBM sample was obtained from excess material collected for clinical purposes from de-identified brain tumor specimens. Nuclei were prepared from this anonymous sample according to Krishnaswami et al. ^43, 44^. Frozen tissue was placed in a Dounce homogenizer containing 1 mL of homogenization buffer (HB: 1 μM DTT, 250 mM sucrose, 25 mM KCl, 5 mM MgCl2, 10 mM tris, 1x protease inhibitor, 0.4 U/μL RNAse, 0.2 U/μL SUPERase In) and subjected to 15 strokes using the loose piston and 15 strokes using the tight piston. The homogenate was divided into 5 volumes and each was strained twice. The filtered homogenate was combined into a tube and centrifuged at 1000 g for 10 minutes at 4 °C. The pellet was resuspended in 1000 μL of HB and strained with 4 strainers (250 μL of homogenate for each strainer). The strained homogenate was again combined into a tube and centrifuged at 1000 g for 10 minutes at 4 °C. The pellet was resuspended in 250 μL of HB to which 250 μL of 50% iodixanol dilution media (IDM: 250 mM sucrose, 150 mM KCl, 30 mM MgCl2, 60 mM tris) was added resulting in a 25% IDM/nuclei suspension. This suspension was carefully layered on top of a tube containing 50% IDM at the bottom and 29% IDM at the surface, and centrifuged at 18,000 g for 23 minutes at 4 C. The nuclei formed a white pellet or sheet in the middle of the tube between the layers. All solution around this pellet was removed – first from the top and then from the bottom, and the pellet was resuspended in 1 mL of chilled PBS and centrifuged (500 g for 5 minutes at 4 °C). The supernatant was removed and the pellet was then treated with DEFND buffer for 5 minutes on ice. Immediately following this incubation, 5 mL of NIB with RNAse inhibitors was added at the nuclei were centrifuged at 500 g for 5 minutes at 4 °C. The pellet was resuspended in 10x Genomics nuclei buffer and a fraction was stained with SYBR green and counted on a Countess with a GFP filter set. Nuclei were then tagmented and prepared for single-cell sequencing using the Chromium Multiome protocol (10x Genomics #1000285, #1000230, #1000215).

### Bulk RNA Library Preparation

Bulk RNA libraries were created for xSDS treated, LAND treated, and untreated BJ fibroblast nuclei with a modified PLATEseq protocol^45, 46^. Following treatment (see DEFNDseq, xSDS, and scATAC protocols above), 5,000 nuclei were lysed in Buffer RLT Plus (QIAGEN #1053393) for each condition. RNA from the lysate was extracted and purified with an RNeasy Mini Kit (QIAGEN #74104) following the manufacturer’s protocol, and eluted in 30 μL of H2O. Of this, 23 μL were mixed with reverse transcription master mix (1.25 μL of 10 μM scc primers, 10 μL Maxima RT buffer (ThermoFisher #EP0753), 5 μL dNTPs (New England Biolabs #N0447L), 2.5 μL SUPERase In, 1.25 μL SMART TSO, 2.5 μL Maxima H Minus Reverse Transcriptase (ThermoFisher #EP0753), and 0.5 μL 10% tween 20 (Millipore Sigma #P9416)). Reverse transcription proceeded on a thermocycler (42 °C for 90 minutes; 10 cycles: 50 °C for 2 minutes, 42 °C for 2 minutes; 75 °C for 10 minutes; 4 °C hold). Then, 1.25 μL of exonuclease I (New England Biolabs #M0293S) and 12.08 μL of H2O were added to the RT product and incubated on a thermocycler (37 °C for 30 minutes, 85 °C for 15 minutes, 75 °C for 30 seconds, 4 °C hold). The product was cleaned up using Dynabeads MyOne Silane (ThermoFisher #37002D). Briefly, silane beads in buffer (30 μL silane beads, 90 μL Buffer RLT Plus, 60 μL EtOH) were mixed with the exonuclease product for 10 minutes, washed twice with 80% EtOH and dried. The cDNA was eluted from the beads with 50 μL of H2O. The cDNA was amplified by combining 25 μL of the silane purified product with PCR master mix (25 μL 2X Kapa HotStart Mix (Kapa Biosystems #KK2602), 2 μL of 5 μM SMART PCR primer) and amplified on a thermocycler (98 °C for 3 minutes; 18 cycles: 98 °C for 20 seconds, 67 °C for 15 seconds, 72 °C for 5 minutes; 72 °C for 5 minutes; 4 °C hold). Following amplification, the product was cleaned up using Ampure XP beads, and analyzed with a BioAnalyzer 2100 using a High Sensitivity DNA Chip (Agilent Technologies #5067-4626)^47^.

### Mixed Species Experiment

U87 and 3T3 cells were dissociated from their culture dishes, counted, and mixed together (1,000,000 cells each). The mixed cells were then treated according to the DEFND-seq protocol. Gene expression and genomic libraries were sequenced separately. Sequencing reads were analyzed according to the snRNA-seq Data Processing and snDNA-seq Data Processing sections below. Cells in the gene expression data were considered human or mouse if they had more than 80% human or mouse purity calculated from gene expression reads. Similarly cells in the gDNA data were considered human or mouse if they had more than 80% human or mouse purity calculated from gDNA fragments.

### Illumina Sequencing

Libraries were sequenced on Illumina sequencers according to 10x Genomics’ suggestions. LAND, xSDS, and scATAC-seq libraries were sequenced on Illumina NextSeq instruments using NextSeq High Output Kit (Illumina #20024907) with 64 cycles for read 1, 64 cycles for read 2, 16 cycles for index 1, and 16 cycles for index 2. These runs were all loaded with 1.5 pM of library. DEFND-seq libraries were sequenced on NextSeq and NovaSeq platforms. The GEX was loaded with 1.8 pM of library and sequenced using a 150 cycle high output kit with 28 cycles for read 1, 102 cycles for read 2, 10 cycles for index 1, and 10 cycles for index 2. The gDNA library was sequenced either using a NextSeq High Output Kit (150 cycles) or a NovaSeq S4. When using NextSeq, 64 cycles were used for read 1, 64 cycles for read 2, 8 cycles for index 1, and 8 dark cycles followed by 16 cycles for index 2. A custom recipe is needed to allow the 8 initial dark cycles on the NextSeq. For the gDNA library on the NovaSeq, 300 pM was loaded and 99 cycles were used for read 1, 99 cycles for read 2, 8 cycles for index 1, and 24 cycles for index 2. All libraries were sequenced with 1% PhiX regardless of the instrument used.

### Element Sequencing

The GBM tumor gDNA library was also sequenced using Element Aviti technology. A fraction (0.5 pmols) of the library from the 10x Genomics protocol was circularized with an Element Adept kit (Element Biosciences #830-00007) following the manufacturer’s protocol. Libraries were quantified with SYBR-based qPCR (ThermoFisher #AB0765). Circularized libraries were denatured and diluted for a target loading concentration of 6 pM with 2% PhiX. The library was loaded on an Aviti 2×150 Sequencing Kit with Adept primers and sequenced with 151 cycles for read 1, 151 cycles for read 2, 8 cycles for index 1, and 24 cycles for read 2. Fastqs were created using Element’s bases2fastq software.

### snRNA-seq Data Processing

snRNA-seq data were processed as described previously^47^. The code is available at (https://github.com/simslab/DropSeqPipeline8). Briefly, we aligned the transcriptomic reads (read 2) to the human or mouse genome and transcriptome annotation using *STAR* v2.7.0d^48^. For human data sets other than the mixed species experiment, we used GRCh38/Gencode v24. For the mixed species data set, we used a concatenated genome and annotation comprised of GRCh38/Gencode v32 and GRCm38/Gencode vM23. Because we are analyzing nuclei that contain a significant amount of unspliced mRNA, we identified reads that uniquely and strand-specifically aligned to the entire gene body of each gene (including exons, introns, intron-exon junctions, and exon-exon junctions), assigning an address to each aligned read containing the gene name, cell-identifying barcode sequence, and unique molecular identifier (UMI) extracted from read 1. Next, we demultiplexed the resulting table, correcting sequencing errors in the barcodes as described in Yuan and Sims^47^ to produce a count matrix for each sample.

### snRNA-seq Clustering and Visualization

For the BJ fibroblasts, we analyzed nuclei with >3,200 unique transcripts and >200,000 unique DNA fragments detected. For the human GBM sample, we analyzed nuclei with >1,000 unique transcripts and >100,000 unique DNA fragments detected. We identified highly variable genes from the snRNA-seq count matrices based on their deviation from the gene drop-out curve as described in Levitin et al.^49^ and used them to construct a Spearman’s correlation matrix and k-nearest neighbor graph from which we performed unsupervised clustering using Louvain community detection as implemented in Phenograph^50^ and visualized with Uniform Manifold Approximation Projection (UMAP)^51^. The cluster-enriched genes that appear in gene expression heatmaps (Figures 2, 4) are selected to highlight certain biological features of each cluster, but are all statistically enriched in a cluster with FDR<0.05 based on the binomial test as described in Shekhar et al.^52^. A computational pipeline for all this analysis is available here (https://github.com/simslab/cluster_diffex2018).

### snDNA-seq Data Processing

snDNA-seq and snATAC-seq data were processed using a custom pipeline available here (https://github.com/simslab/dna10x). The pipeline attempts to mimic some of the basic data processing procedures that are implemented in the 10x Genomics Cell Ranger software packages for analysis of snATAC-seq and Multiome data, including the formatting of output files. We generated sample-demultiplexed fastq files using the *mkfastq* command in *cellranger-atac* v2.0.0 (10x Genomics). We aligned paired-end reads 1 and 2 to the human or mouse genome using *bwa mem* v0.7.17-r1188^53^ after removing a standard adapter sequence (CTGTCTCTTATACACATCT) from each read using *cutadapt* v2.8^54^. We then extracted all reads with an alignment score that was >90% of the read length and insert size <1 kb. We also implemented an error correction procedure for the cell-identifying barcodes from read 3. Briefly, for cell-identifying barcodes that do not appear in the standard list provided by 10x Genomics, we first determine the frequency *f_alt_* of each barcode in the standard list in the data set. For each observed barcode sequence that has a Hamming distance of one from a sequence on the standard list, we estimate the posterior probability *p_alt_* that the barcode deviates from the standard list due to sequencing error using the quality scores for the putatively errant base provided by Illumina or Element *p_alt_* = *f_alt_10^-q/10^* where *q* is the quality score. If *p_alt_*>0.975 for any alternative barcodes, then we replace the errant barcode with the alternative barcode from the standard list with the highest value of *p_alt_*. Next, we establish read addresses for each alignment that passes the filter described above comprised of a cell-identifying barcode, chromosome, a fragment start position, a fragment end position and collapse identical fragments. Finally, we sort the fragments by alignment position on the genome and output a table in the format of the “fragments.tsv” file typically produced by Cell Ranger.

### snDNA-seq Data Analysis and Visualization

The data processing pipeline described above produces output files that are formatted similarly to those of Cell Ranger, thus the data can be analyzed by multiple software packages that have been developed for snATAC-seq or joint snRNA/ATAC-seq data. We analyzed the data using a recently developed Python/Rust implementation of SnapATAC, called SnapATAC2^55, 56^. Specifically, we used the *pp.import_data* function to convert the standard “fragments.tsv” file from our data processing pipeline into the h5ad format. This function also computes basic features of each profiled nucleus including the number of unique fragments detected, the transcription start site (TSS) enrichment score, and the duplication rate. We used the *pp.make_tile_matrix* function to produce all of the copy number variation (CNV) plots and heatmaps. The function bins the unique fragments detected along each chromosome in each nucleus for a given bin size (0.1 MB and 1 MB used for the figures shown here as indicated).

Figs. 2, 3, 4 and 5 show CNV plots, heatmaps, and corrected coefficients-of-variation (CVs) for individual nuclei or clusters. The procedures used to correct coverage bias are sample-dependent. Ideally, for a given sample, we would identify a high-confidence subpopulation of euploid cells and normalize all cells by their median binned CNV profile. This would significantly correct for multiple sources of coverage bias that are present in all nuclei (e.g. GC content bias, Tn5 insertion bias). This is exactly what we did for the BJ fibroblast data as shown in Fig. 3. First, we identified any problematic bins using the *gcCounter* and *mapCounter* functions in HMM Copy / HMM Copy Utils^57, 58^. Specifically, we eliminated any bins where *gcCounter* did not output a GC content or where *mapCounter* scores the mappability of a bin as <0.95. Next, we identified the top 50 least noisy BJ fibroblast nuclear profiles by computing the CV in coverage (number of unique fragments per bin after normalizing by the total number of unique fragments) of all somatic chromosomal bins that were not eliminated by *gcCounter* or *mapCounter*. Finally, we computed the median coverage profile of these top 50 profiles, multiplied these values by two for all sex chromosomal bins, eliminated any bins with a median coverage of zero for the top 50 profiles, and normalized all nuclear profiles by the resultant low-noise profile. For each nuclear profile, we divided the resulting, corrected coverage profile by the median corrected coverage of all somatic chromosomal bins and multiplied by two to arrive at an estimated copy number for each bin.

For the GBM sample, there were too few euploid cells to take the approach described above. As an alternative, we reduced the GC-bias in our copy number profiles by directly correcting for it. We computed the GC content of each 100 kb genomic bin using *gcCounter* and the median coverage of each bin. We then median-filtered the coverage as function of GC content for all bins with GC content between 0.3 and 0.6 with a window-size of 0.001 as plotted in Supplementary Information, Figure S6. Finally, we fit a correction curve to this plot using LOWESS regression as implemented in the Python *statsmodels* function *lowess* with default parameters. For each 100 kb bin, we computed an interpolated correction value from the LOWESS regression fit by which we normalized the coverage profile of each nucleus. Similar to the BJ fibroblast data, we eliminated any bins with mappability <0.95 as well as any bins with GC content <0.3 or >0.6. We divided the resulting corrected coverage profiles by the median corrected coverage for Chr. 2 and multiplied by two to obtain an estimated copy number for each bin in each nucleus. We chose Chr. 2 because it is the largest somatic chromosome with a copy number that was uniformly double that of the corresponding sex chromosomes both in this dataset in a previously published bulk WGS profile of this same tumor^23^ (i.e. Chr. 2 is likely diploid, even in the transformed glioma cells).

We attempted to identify somatic single nucleotide variants (SNVs) from the DEFND-seq genomic data for the GBM sample. This was challenging because a germline genome of the subject was unavailable. We used the Genome Analysis Toolkit (GATK)^59^ tumor-only pipeline for this analysis including removal of duplicate reads from the original bam file produced by *bwa mem* with the Picard *gatk MarkDuplicates* command, base quality score recalibration with *gatk ApplyBQSR*, and read sorting with *samtools sort*. We called mutations using Mutect2 via the *gatk Mutect2* command with Gnomad germline resource^60^ af-only-gnomad.vcf.gz and the 1000-genomes panel-of-normals 1000g_pon.hg38.vcf.gz provided GATK. This analysis cross-references mutations detected by Mutect2 with databases of common SNPs, marking them as potentially originating from the germline. For further filtering, we used *gatk GetPileupSummaries* and *gatk CalculateContamination* to generate a contamination table for filtering the Mutect2 variant calling format (VCF) file with *gatk FilterMutectCalls*. Finally, we used the *table_annovar.pl* Perl script as part of the Annovar^61^ package to annotate the resulting filtered VCF file.

We performed the analysis described above to produce annotated VCF files for the GBM sample originating from two different sequencing runs – one obtained using an Illumina NovaSeq 6000 and the other with an Element Aviti sequencer. We reasoned that these two platforms would have somewhat different biases and error profiles, which could result in increased variant calling accuracy. We first intersected the two VCF files using *bcftools*^62^, identifying variants found in independent analyses from both sequencers. We then cross-referenced the intersected VCF file with the Catalogue of Somatic Mutations in Cancer (COSMIC) database^29^, which revealed a missense variant in PREX1 that had been previously confirmed as a somatic SNV in both a pancreatic ductal adenocarcinoma case and another case of GBM. Regions containing SNVs were visualized with the Integrated Genomics Viewer^63^ and presented in Figure 5.

### Custom Oligos

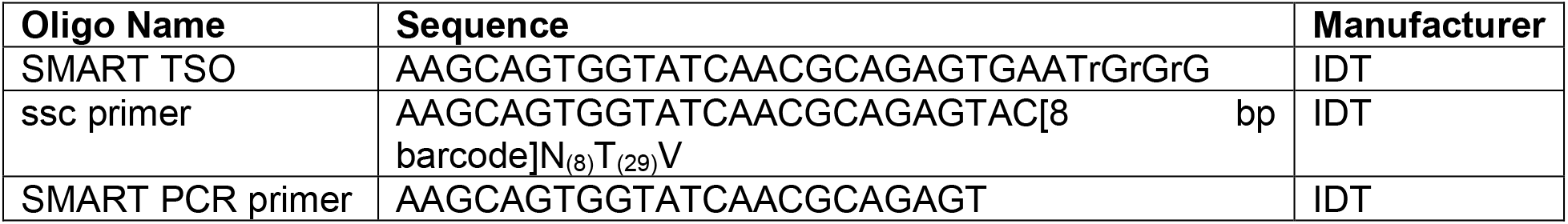

## Data Availability

The sequencing data and count matrices reported in this paper are available in the following database: Gene Expression Omnibus GSE224149: https://www.ncbi.nlm.nih.gov/geo/query/acc.cgi?&acc=GSE224149

## Code Availability

All source code is available on the Sims Lab GitHub repository, https://github.com/simslab/, including code for generating expression count matrices (DropSeqPipeline8), clustering and differential expression (cluster_diffex2018), and genomic DNA analysis (dna10x).

## Acknowledgements

P.A.S. and S.Z. were supported by a RISE grant from Columbia University. S.Z. was supported by NIH/NCI grant R01CA275184. P.T. was supported by the 1.1. Rabi Scholars Program of Columbia University. P.A.S. was supported by NIH/NCI grant U54CA209997. P.A.S., P.C., and J.N.B. were supported by NIH/NINDS grant R01NS103473.

## Author Information

### Affiliations

**Department of Systems Biology, Columbia University Irving Medical Center, New York, NY**

Timothy R. Olsen, Pranay Talla, & Peter A. Sims

**Department of Neurological Surgery, Columbia University Irving Medical Center, New York, NY**

Julia Furnari & Jeffrey N. Bruce

**Department of Pathology & Cell Biology, Columbia University Irving Medical Center, New York, NY**

Peter Canoll & Shan Zha

**Institute for Cancer Genetics, Columbia University Irving Medical Center, New York, NY**

Shan Zha

**Department of Pediatrics, Columbia University Irving Medical Center, New York, NY**

Shan Zha

**Sulzberger Columbia Genome Center, Columbia University Irving Medical Center, New York, NY**

Peter A. Sims

**Department of Biochemistry & Molecular Biophysics, Columbia University Irving Medical Center, New York, NY**

Peter A. Sims

### Contributions

T.R.O. and P.A.S. conceived experiments. T.R.O., P.T., and P.A.S. analyzed data. T.R.O. and P.T. performed the experiments. J.F. managed biobanked clinical samples. P.C. and J.N.B. procured and characterized clinical specimens. T.R.O. and P.A.S. wrote the manuscript. All authors reviewed and edited the manuscript.

## Ethics Declarations

For studies of human specimens, all tissue was procured from de-identified patients who provided written informed consent through a protocol approved by the Columbia University Institutional Review Board (IRB-AAAJ6163).

---

**Extended Data Figure 1:**
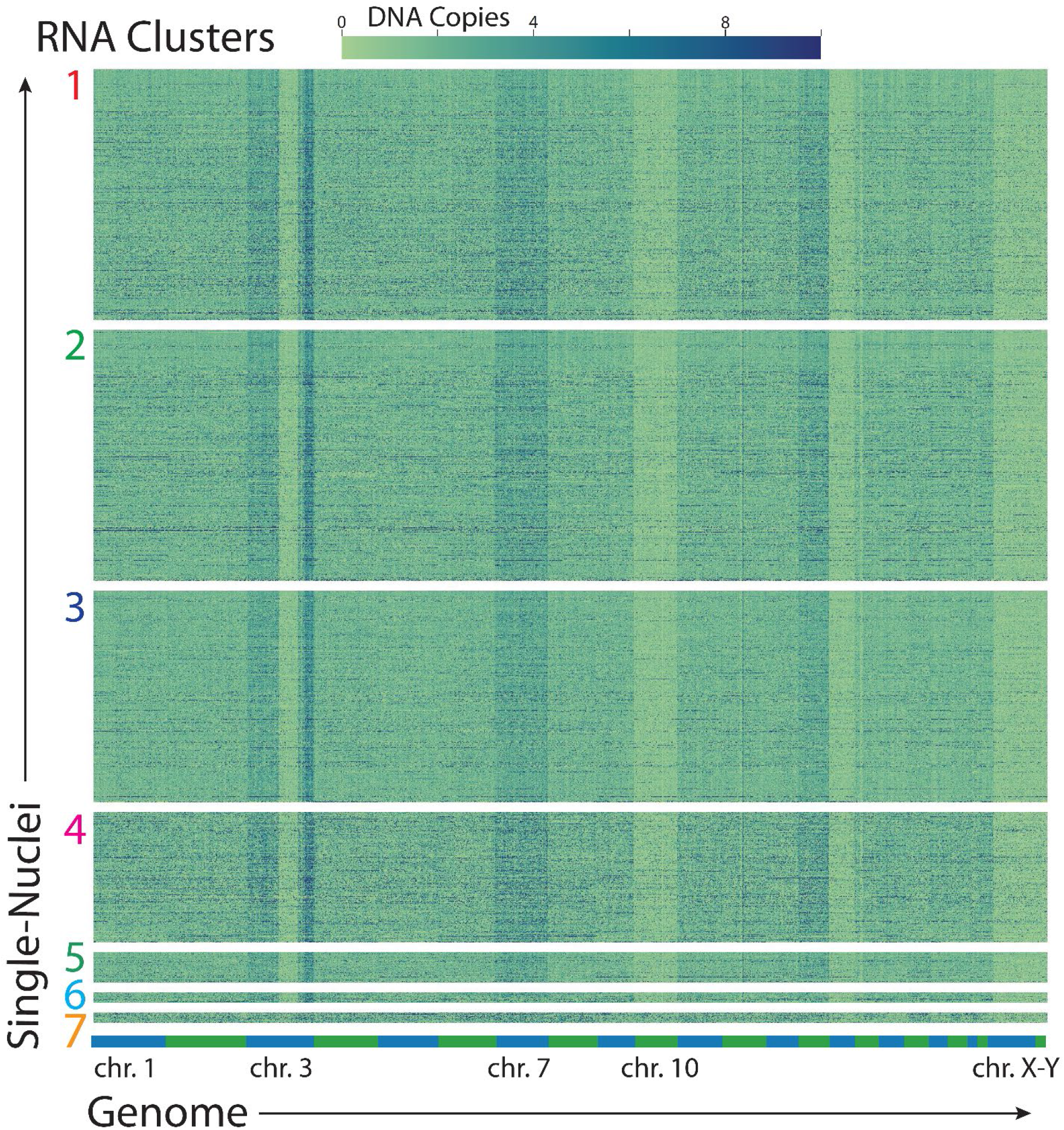
Copy number variation heatmap of GBM tumor cells. Cells are grouped by gene expression cluster. Chromosome location reference on bottom margin.

