## Supplementary Figures 1-6 for "Scalable co-sequencing of RNA and DNA from individual nuclei"

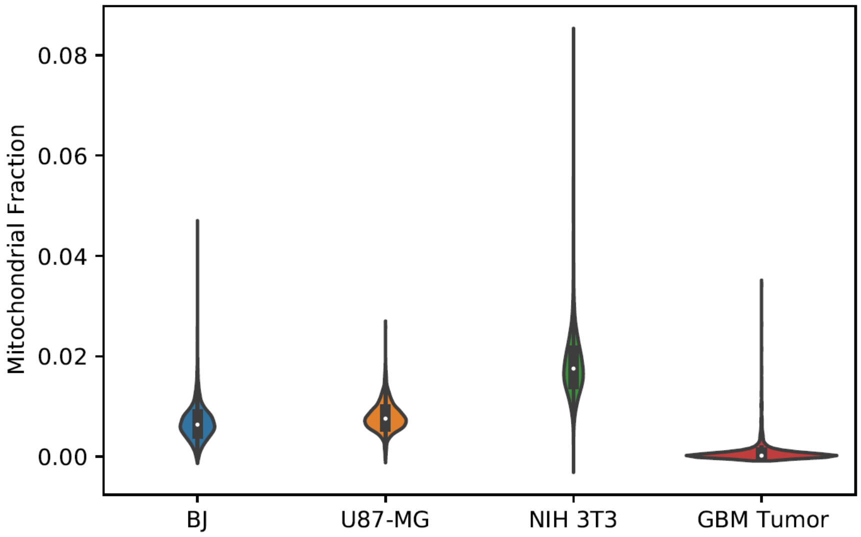


**Figure S1**: DEFND-seq mitochondrial alignment rates for gene expression reads for various cell types.

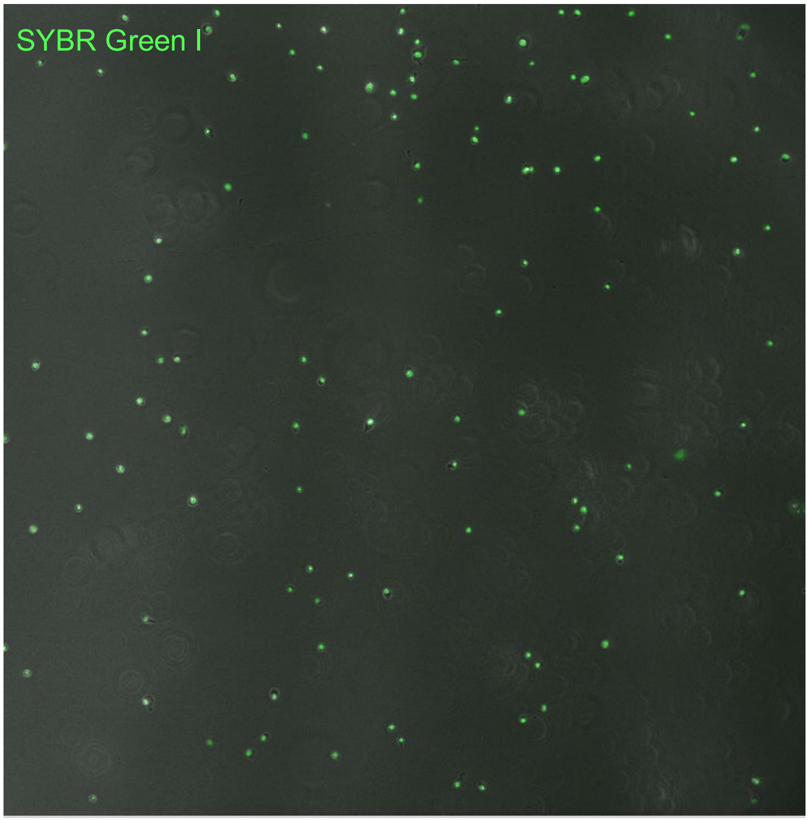


**Figure S2**: Fluorescence microscopy image of SYBR Green I stained 3T3 and U87 nuclei that have been nucleosome depleted with lithium diiodosalicylate prior to tagmentation.

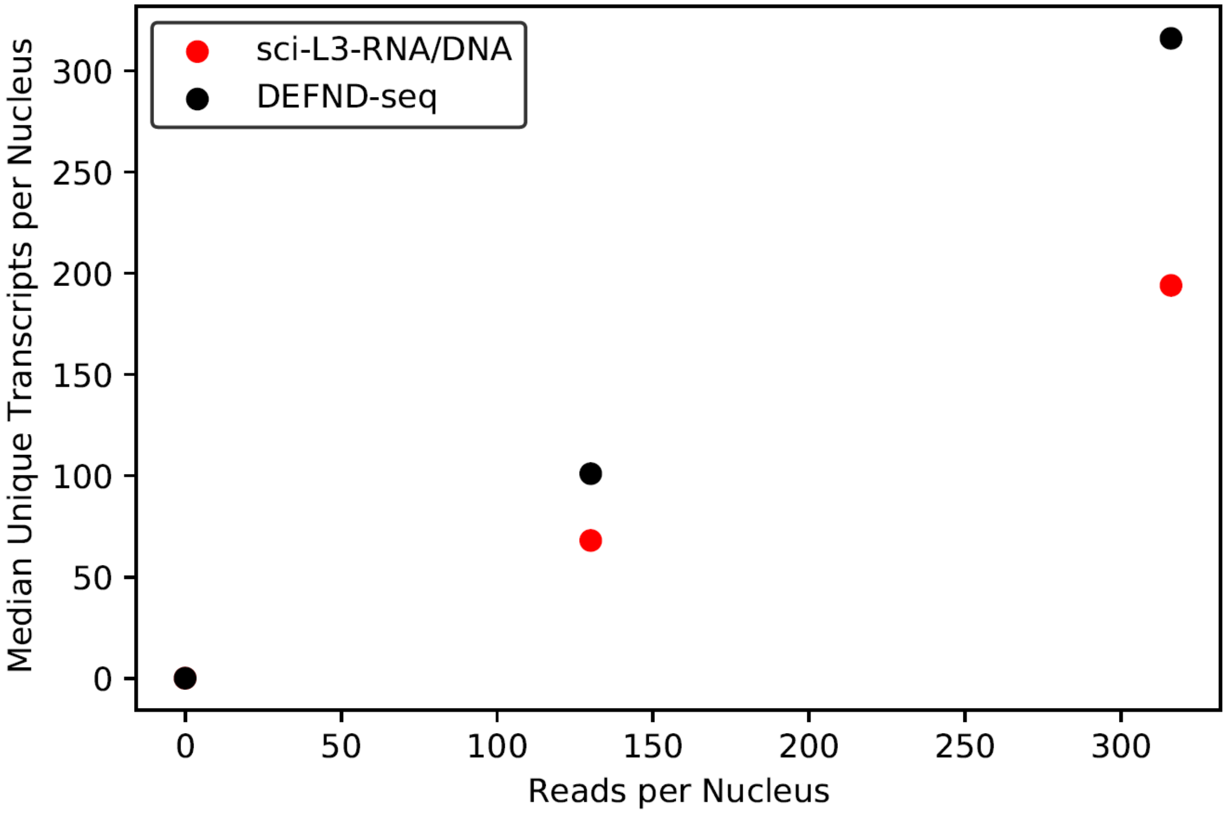


**Figure S3**: Median unique transcripts per nucleus for RNA/DNA co-assays at various sequencing depths.

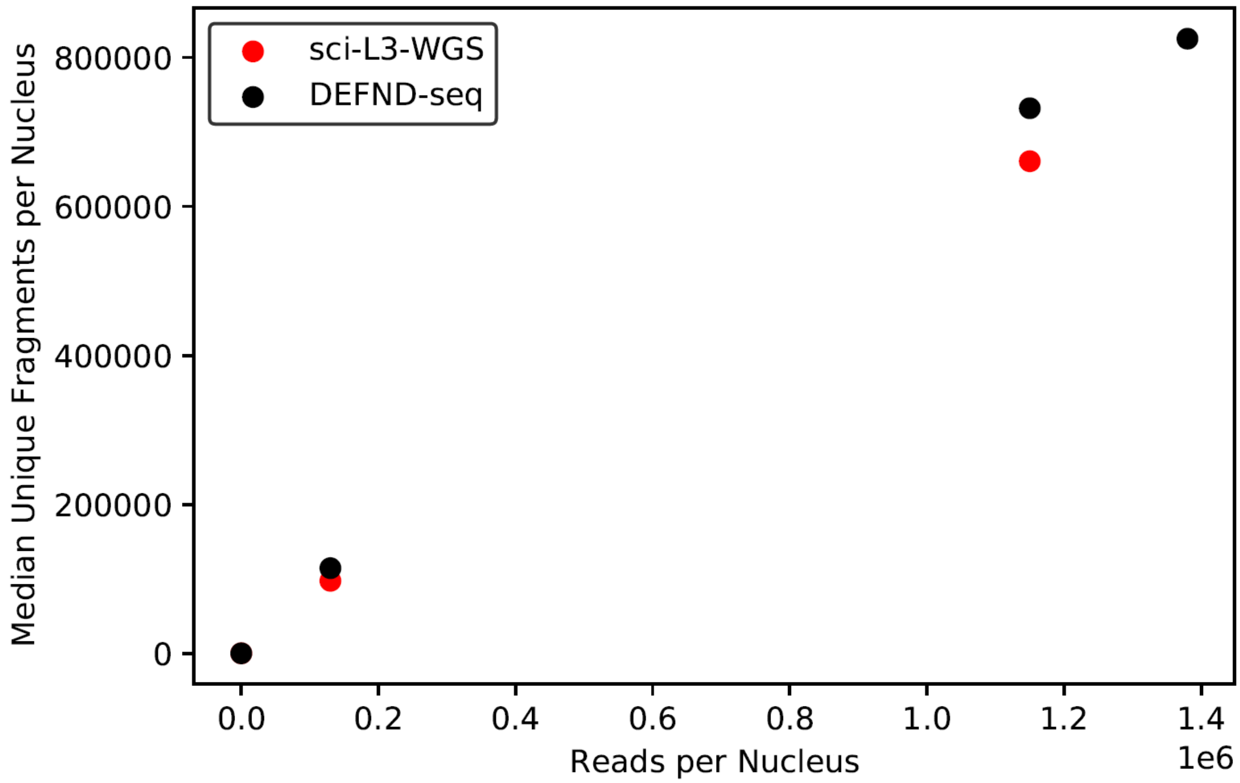


**Figure S4**: Median unique fragments per nucleus for RNA/DNA co-assay DEFND-seq and gDNA-specific assay sci-L3-WGS at various sequencing depths.

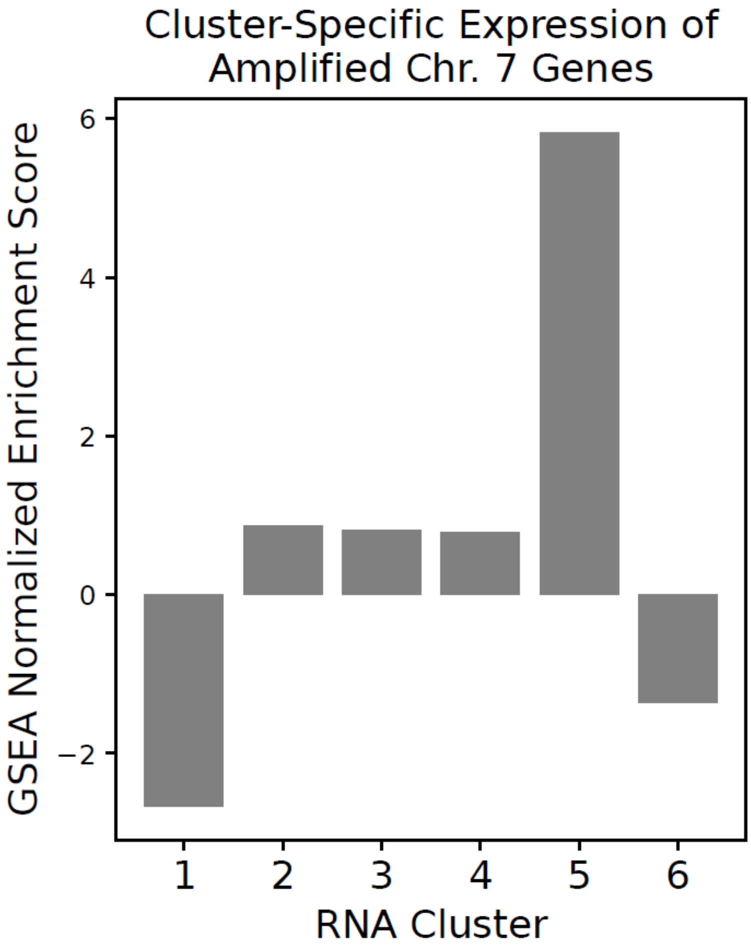


**Figure S5**: Gene set enrichment analysis (GSEA) for expression of genes that are amplified in Chr. 7q region. The GSEA score is calculated for each gene expression cluster. Cluster 5 cells are enriched in the 7q amplification and correspondingly express higher levels of the amplified genes.

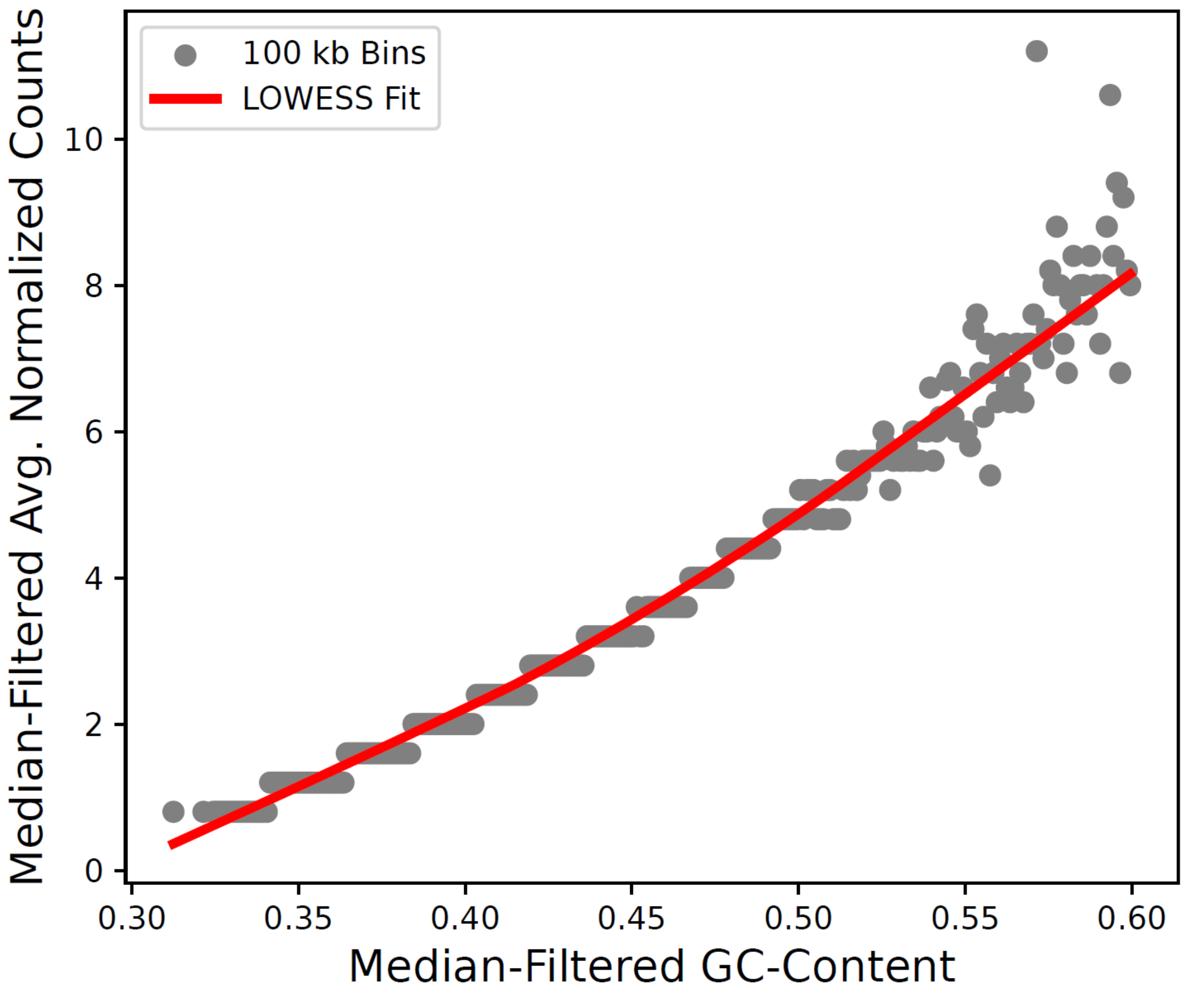


**Figure S6**: GC content correction of GBM coverage. Window size of 0.001 for GC content.
